# Acquisition of the Spindle Assembly Checkpoint and its modulation by cell fate and cell size in a chordate embryo

**DOI:** 10.1101/2022.05.18.492463

**Authors:** Marianne Roca, Lydia Besnardeau, Elisabeth Christians, Alex McDougall, Janet Chenevert, Stefania Castagnetti

## Abstract

The spindle assembly checkpoint (SAC) is a surveillance system which preserves genome integrity by delaying anaphase onset until all chromosomes are correctly attached to spindle microtubules. Recruitment of SAC proteins to unattached kinetochores generates an inhibitory signal that prolongs mitotic duration. Chordate embryos are atypical in that spindle defects do not delay mitotic progression during early development, implying that either the SAC is inactive or the cell-cycle target machinery unresponsive. Here we show that in embryos of the chordate *Phallusia mammillata* the SAC delays mitotic progression from the 8^th^ cleavage divisions. Unattached kinetochores are not recognized by the SAC machinery until the 7^th^ cell cycle when the SAC is acquired. Following acquisition, SAC strength, manifest as the degree of mitotic lengthening induced by spindle perturbations, is specific to different cell types and is modulated by cell size showing similarity to SAC control in early *Caenorhabditis elegans* embryos. We conclude that SAC acquisition is a process likely specific to chordate embryos, while modulation of SAC efficiency in SAC proficient stages depends on cell fate and cell size similarly to non-chordate embryos.

## Introduction

The spindle assembly checkpoint (SAC) operates during mitosis to ensure fidelity of chromosome segregation and loss of SAC function is often associated with chromosome segregation errors and the generation of daughter cells with an abnormal chromosome number, a condition called aneuploidy, deleterious for cell viability and organismal survival (Chunduri and Storchová, 2019). The SAC monitors kinetochore-spindle microtubule attachments through the recruitment of SAC components to unattached kinetochores until all kinetochores have acquired proper bipolar attachments to spindle microtubules (Jia et al., 2013; Musacchio and Salmon, 2007). Kinetochore localization of SAC proteins promotes the formation of a soluble complex, the mitotic checkpoint complex, which by sequestering Cdc20 acts as a potent inhibitor of APC/C (an E3 ubiquitin ligase required for anaphase onset and mitotic exit) delaying chromosome segregation and mitotic progression (De Antoni et al., 2005; Sudakin et al., 2001).

Even though the SAC is essential to preserve ploidy, the response to spindle defects varies across cell types, and early embryos of several species dispense with the SAC (Chenevert et al., 2020). This variability in SAC efficiency has been attributed to differences in cell size and cell identity. However, the available results are contradictory. Studies carried out in *C. elegans* embryos showed that the SAC becomes more efficient, delaying mitosis for longer, at each division following fertilization, as cell volume is reduced (Galli and Morgan, 2016; Gerhold et al., 2018). These data can be explained by a modulation model whereby SAC signal generated at kinetochores is diluted in the cytoplasm and the strength of SAC response is influenced by kinetochore-to-cytoplasmic ratio. Consistent with this notion, in *Xenopus* egg extracts, the mitotic checkpoint can be activated by increasing the concentration of added sperm heads (Minshull et al., 1994) and in mouse reduction of oocyte size results in a stronger SAC response during meiosis (Kyogoku and Kitajima, 2017; Lane et al., 2017). Crucially however, artificially changing cell volume does not affect the timing of SAC acquisition either in frog (Clute and Masui, 1995; Clute and Masui, 1997) or in fish embryos (Zhang et al., 2015), suggesting that modulation of SAC strength and SAC acquisition might be two different phenomena.

It has proven difficult to tease apart SAC acquisition from SAC modulation. However, a prerequisite of the modulation hypothesis is that SAC proteins do accumulate on unattached kinetochores but the inhibitory signal produced is too weak to induce a delay in mitotic progression. In SAC-proficient *C. elegans* embryos, Mad1 and Mad2 localize to unattached kinetochores already from the 2-cell stage (Galli and Morgan 2016). Kinetochore localization of SAC components has not been analysed in SAC-deficient fish and frog embryos. However, in embryos of the ascidian *Phallusia mammillata* Mps1, Mad2 and Mad1 do not accumulate to unattached kinetochores in 2-cell stage embryos (Chenevert et al., 2020).

Here we find that in *P. mammillata* embryos the SAC becomes efficient at the 8^th^ cell cycle. The appearance of a functional SAC correlates with SAC acquisition rather than SAC modulation since recruitment of SAC proteins to unattached kinetochores first occurs at the 7^th^ cell cycle. We show that, following SAC acquisition SAC strength increases at each cell division and is influenced by cell identity, similarly to SAC proficient non-chordate embryos (Gerhold et al., 2018). We suggest that within same-fated cells SAC strength is then modulated by changes in cell size. In the ventral ectoderm, the difference in SAC activity observed between anterior and posterior cells requires GSK3 activity to enhance SAC strength in anterior cells. Although SAC acquisition appears to be specific to chordate embryos (Chenevert et al., 2020), SAC modulation in chordates depends on cell fate and cell size similarly to non-chordate embryos.

## Results and Discussion

We previously showed that the SAC is inefficient in *P. mammillata* 2-cell stage embryos where nocodazole treatment prevents spindle formation, but induces only a limited mitotic delay (Chenevert et al., 2020). To determine when the SAC becomes functional in *P. mammillata* embryos, we analyzed mitotic duration in the absence of spindle microtubules (Fig. S1), from the 2^nd^ to 10^th^ cell cycle following fertilization. Mitotic duration was measured as the time spent between nuclear envelope breakdown (NEB), and nuclear envelope reformation (NER; Fig. 1A). Consistent with earlier studies (Dumollard et al., 2017), mitotic duration in control embryos (DMSO) was constant at all analyzed stages (Fig. 1B) and lasted on average 10 minutes. As for 2-cell stage embryos (Chenevert et al., 2020), mitotic duration was briefly extended in 4- to 64-cell stage embryos treated with nocodazole, compared to DMSO-treated embryos (1.5-fold) (Fig. 1B). From the 8^th^ cell cycle, however, in nocodazole-treated embryos mitotic duration was prolonged more extensively at each cell cycle, extending by 2.5-fold (24.5±8.5 minutes) at the 8^th^ cycle (128-cell), by 4.5-fold (37.3±14.5 minutes) at the 9^th^ cycle (256-cells) and by 6.5-fold (64.3±20 minutes) at the 10^th^ cycle (512-cells). To determine whether this mitotic delay required SAC activity, we then measured mitotic duration in the presence of nocodazole in SAC impaired embryos (Fig. 2A). Mutations of three phosphorylation sites in the human SAC protein Mad2 produce a dominant negative form of Mad2 (Mad2-DN) that impairs SAC activity (Fig. S2A) (Wassmann et al., 2003a; Wassmann et al., 2003b). We tested the effect of Mad2-DN on mitotic duration during *P. mammillata* development. At the 2-cell stage, overexpression of Mad2-DN did not affect mitotic duration, either in the presence (Fig. S2B) or in the absence (Fig. 2B) of microtubules. However, both at the 8^th^ and the 9^th^ cell cycles mitotic duration in the presence of nocodazole was significantly reduced by Mad2-DN overexpression (Fig. 2B) lasting only 1.5 times longer in nocodazole-treated than in DMSO-treated embryos (128-cell: DMSO = 9.8±1.6 to nocodazole = 15.6±3.5 minutes; 256-cell: 9.9±1.8 to 13.6±2.4 minutes), comparable to mitotic duration in nocodazole treated wild-type early embryos (2- to 64-cells). Similarly impairing SAC activity with reversine, a specific inhibitor of Mps1, the kinase that initiates the SAC signaling cascade (Fig. 2A)(Santaguida et al., 2010), prevents mitotic lengthening following nocodazole treatment at the 9^th^ cell cycle (Fig. 2B). Hence, the SAC is required to delay mitotic exit from the 8^th^ cell cycle. During the first seven cell division cycles the SAC could either be inactive, or active but unable to induce a prolonged delay in mitotic duration.

**Figure 1:**
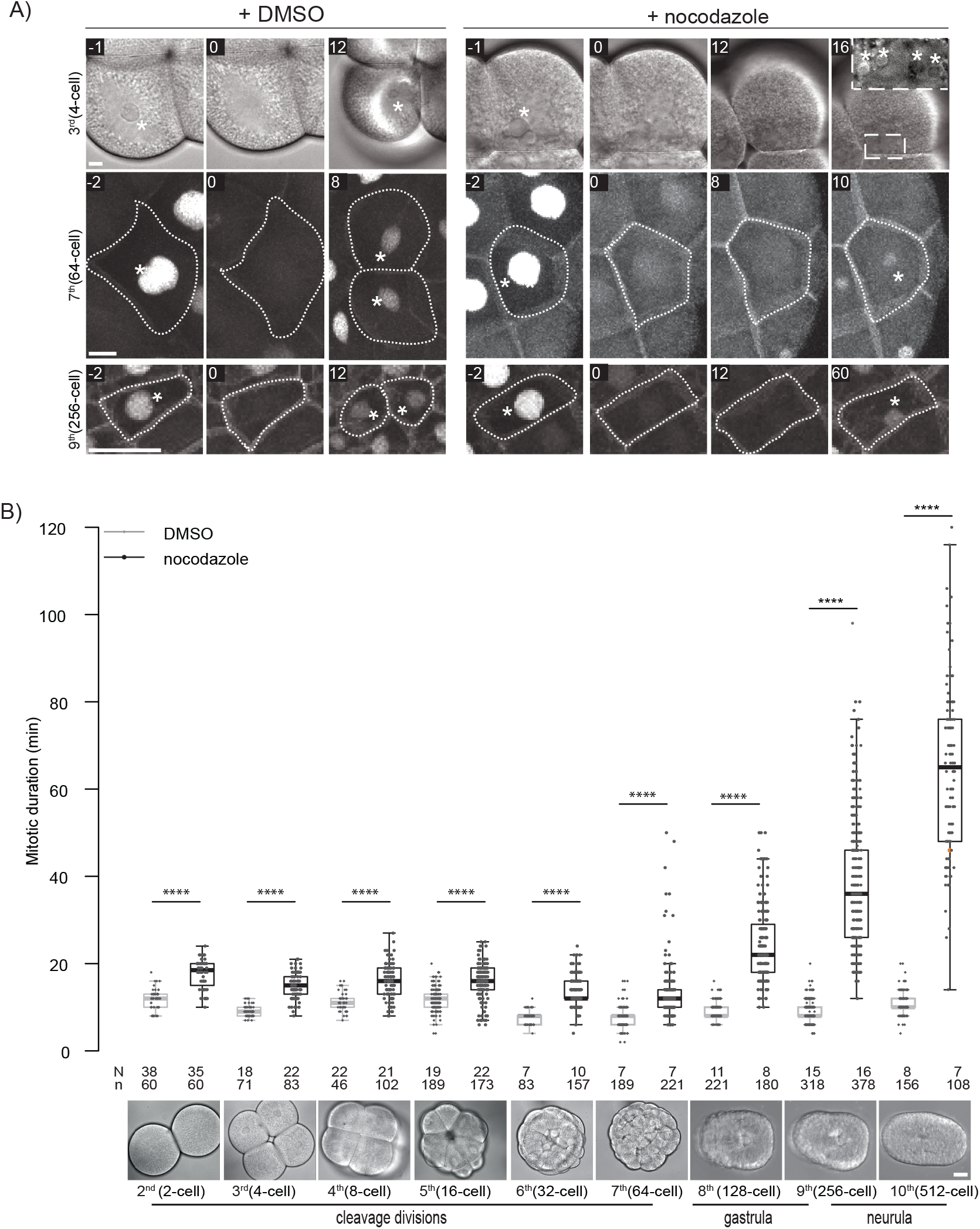
Nocodazole treatment induces a prolonged delay in mitotic progression from the 8^th^ cell cycle. **A)** Representative images of cells from 4- (top), 64- (middle) and 256-cell (bottom) embryos treated with DMSO (left) or with nocodazole (right). Asterisks indicate nuclei. NEB is considered as time 0 and time before or after NEB (minutes) is indicated on each image. See also Movies 1 and 2. Dotted lines indicate cell periphery. Fluorescence images are projections of the confocal Z-stack covering the whole cell. **B)** Quantification of mitotic duration in DMSO (grey) or nocodazole (black) treated embryos from 2^nd^ to 10^th^ cell cycle (2- to 512-cell). Representative brightfield images of *P. mammillata* embryos are presented below the plot for all analyzed stages. Each dot represents a cell. Boxes represent 25-75^th^ percentiles, and the median is shown. Number of analyzed embryos (N) and cells (n) is given for each stage. Test of Wilcoxon-Mann-Whitney performed using all cells: non-significant (ns), p-value ≤ 0.05 (*), ≤ 0.01 (**), ≤ 0.001 (***), ≤ 0.0001 (****). Scale bars are 10 µm in A and 30 µm in B.

**Figure 2:**
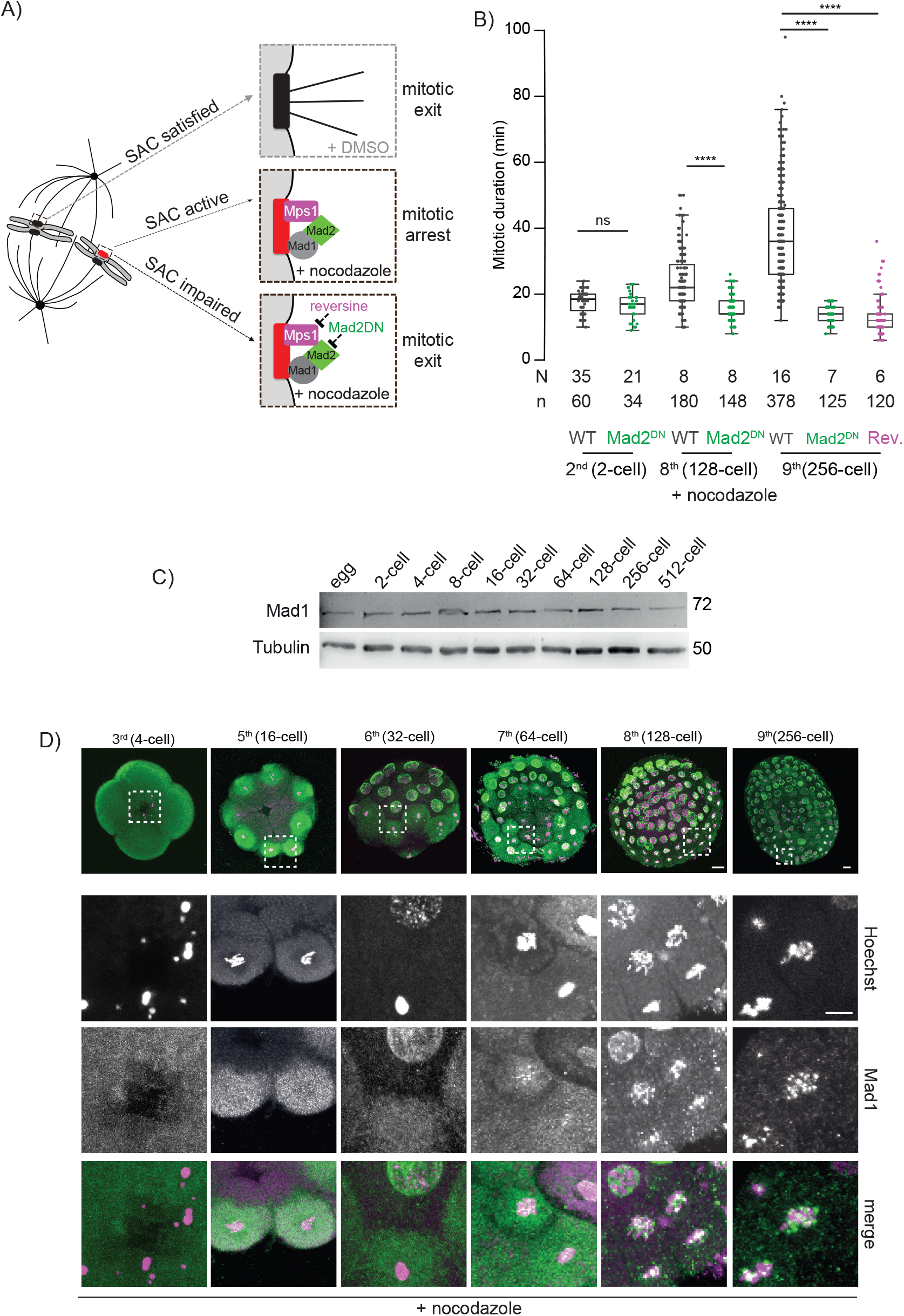
SAC activity is acquired at the 8^th^ cell cycle. **A)** Schematic representation of mitotic chromosomes and their association with spindle microtubules. SAC proteins Mps1, Mad1 and Mad2 localize to an unattached kinetochore, leading to SAC activation and inhibition of APC/C activity (top). Treatment with the Mps1 inhibitor reversine or overexpression of Mad2-DN impairs SAC activity resulting in premature mitotic exit (bottom). **B)** Quantification of mitotic duration in nocodazole treated embryos at 2-, 128- and 256-cell stage in wild-type embryos (black) or following SAC impairment by either overexpression of Mad2-DN (green) or treatment with reversine (magenta). Box plots are as in Figure 1B. See also Movies 3 and 4. **C)** Western blot analysis of Mad1 protein (top) throughout development (egg to 512-cell). Tubulin is used as loading control (bottom). **D)** Representative images of 4-, 16-, 32-, 64-, 128- and 256-cell (left to right) embryos treated with nocodazole, stained with Hoechst (DNA, magenta) and for Mad1 (green). White squares indicate mitotic cells enlarged underneath each image. See also Figure S2. Scale bars are 10 µm.

To distinguish between these two hypotheses, we assessed whether a SAC signal was produced during the early division cycles. As kinetochore localization of SAC components is an essential prerequisite for SAC signaling (Musacchio and Salmon, 2007), we analyzed the presence of Mad1 throughout development and its localization to unattached kinetochores, before and after SAC acquisition, using a specific antibody against *P. mammillata* Mad1 (Chenevert et al., 2020). We find that Mad1 protein is present at similar levels throughout development (Fig. 2C) irrespective of SAC activity. As for yeast and somatic cells (Rodriguez-Bravo et al., 2014), Mad1 localizes to the nuclear envelope during interphase at all developmental stages independently of microtubules (Fig. 2D and S2C). However, as for 2-cell stage embryos (Chenevert et al., 2020), in the presence of nocodazole, Mad1 does not accumulate to mitotic chromosomes in 4- to 32-cell embryos (Fig. 2D). Mad1 foci associated with mitotic chromosomes were observed in 128- and 256-cell stage embryos (53% and 74% of mitotic cells respectively). We also observed weak accumulation of Mad1 on unattached chromosomes in 14% of mitotic cells at the 64-cell stage (Fig. 2D). It is thus possible to distinguish two phases during *P. mammillata* development with respect to SAC activity; first, a SAC-deficient phase when the presence of unattached kinetochores goes undetected by the cell, which continues cycling irrespective of the lack of spindle-kinetochore interactions, and second a SAC-proficient phase when the SAC machinery is recruited to unattached kinetochores, an essential step in the generation of the inhibitory signal that delays mitotic progression until the formation of correct bipolar interactions.

After the 8^th^ cell cycle, SAC response increases both in strength and variability at each cell cycle (Fig. 1B). At this stage, most cells in the ascidian embryo are fate-restricted. We then asked whether SAC strength is a property of different cell identities and analyzed SAC strength in different cell populations. We observed that at the 9^th^ cell cycle, in the presence of nocodazole, cells of the anterior ventral ectoderm spent significantly longer in mitosis than cells of the posterior ventral ectoderm, dorsal ectoderm or notochord (Fig. 3A, B).

**Figure 3:**
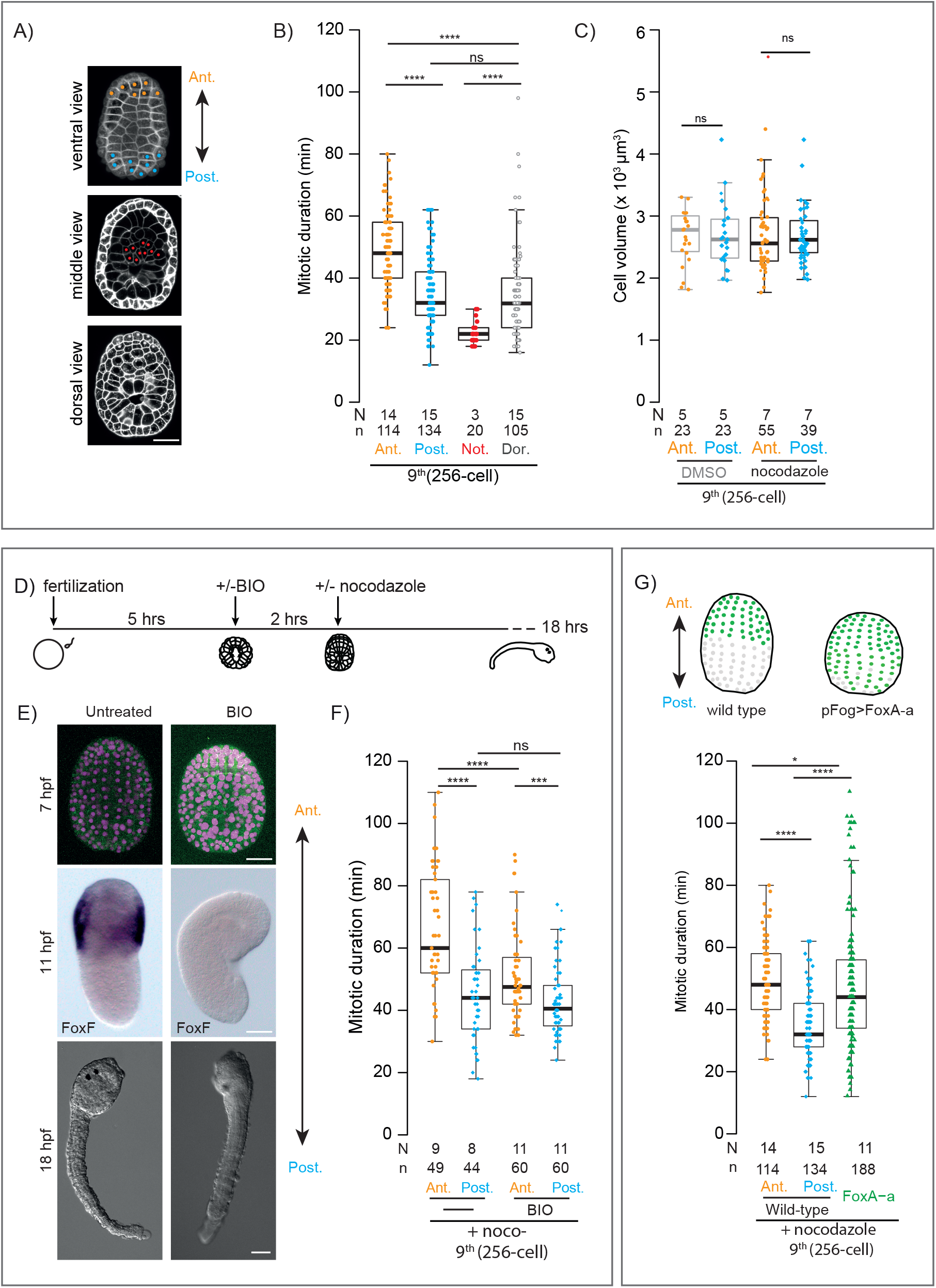
SAC is stronger in anterior ectodermal cells. **A**) Ventral, middle and dorsal views of cell mask stained 256-cell stage embryo, indicating areas where mitotic duration was analyzed: ventral anterior (orange) and posterior (blue) ectoderm, notochord (red) and dorsal ectoderm (grey). **B)** Mitotic duration in ventral anterior ectoderm (orange), ventral posterior ectoderm (blue), notochord (red) and dorsal cells (grey) of nocodazole treated wild type 256-cell stage embryos. **C)** Cell volume of ventral anterior (orange) and posterior (blue) ectodermal cells in 256-cell stage embryos in the presence of either DMSO (grey) or nocodazole (black). **D)** Schematic representation of experiment to analyze impact of GSK3 activity on mitotic duration. **E)** Representative images of control (left) and BIO-treated (right) 256-cell stage embryos (7 hours post fertilization, hpf), expressing NLS-Tomato (magenta) and PH-GFP (green), early tailbud embryos (11hpf) showing the expression pattern of the anterior marker *FoxF* and tadpoles (18 hpf). **F)** Mitotic duration in ventral anterior (orange) and posterior (blue) ectodermal cells of control (DMSO) embryos and BIO-treated embryos at the 9^th^ cell cycle in the presence of nocodazole. Box plot in C is as in Figure 1B. **G)** Top: schematic representation of FoxA-a expression (green) in wild type and pFog>FoxA-a over-expressing embryos. Bottom: mitotic duration in ventral anterior ectoderm (orange), ventral posterior ectoderm (blue) and pFog>FoxA-a overexpressing embryos (green) at the 9^th^ cell cycle in the presence of nocodazole. Scale bars are 30µm.

To dissect what influences this difference in SAC strength we focused on the ventral ectoderm as those cells are easily accessible on the surface of the embryo, and two SAC behaviors can be observed in the same field of view. We first asked whether the difference between these two populations could be explained by a difference in cell size. Using the membrane marker PH-GFP, we measured cell volume and observed a similar range of cell volumes for anterior and posterior ventral ectodermal cells (Fig. 3C), excluding that the difference in mitotic response between these two cell populations is due to a difference in cell size.

We then asked whether the difference in SAC strength along the antero-posterior axis is a consequence of the difference in cell fates. We therefore interfered with patterning along the anterior-posterior axis and analyzed SAC strength at the 9^th^ cell cycle. In the ascidian *Ciona intestinalis*, GSK3 is active in anterior cells and its inhibition from the onset of gastrulation results in loss of anterior identity in the ectoderm with extension of the posterior domain (Feinberg et al., 2019). We observed a similar effect in *P. mammillata* embryos treated with the specific GSK3 inhibitor BIO from the onset of gastrulation (Fig. 3D). BIO-treated embryos did not express the anterior markers *FoxF* (Fig. 3E) (Feinberg et al., 2019)) and *sFRP1/5* (Fig. S3A) (Lamy et al., 2006), and developed into tadpoles with diminished anterior structures (Fig. 3E), consistent with the acquisition of posterior identity in the anterior ventral ectoderm. At the 256-cell stage, BIO-treated embryos maintained their overall shape, allowing orientation of the embryo along the anterior-posterior axis (Fig. 3E). At this stage, mitotic duration was slightly extended in posterior ectodermal cells by BIO treatment alone, but no effect was observed in anterior ectodermal cells (Fig. S3B; wild-type: anterior 11.8±2.9 minutes, posterior 10.7±1 minutes; BIO: anterior 12.1±2.9 minutes, posterior 13.6±3.2 minutes). In the presence of nocodazole, however, (Fig. 3F) mitotic duration was significantly shorter in anterior cells of the ventral ectoderm in BIO-treated embryos than in untreated embryos (untreated: 66.4±19.1 minutes, BIO: 50.9±13.9 minutes), while no difference was observed in ventral posterior ectodermal cells (untreated: 44.7±15.2 minutes, BIO: 43.1±11.3 minutes).

As posteriorization of anterior cells results in reduction of SAC strength we then asked whether anteriorization of the posterior ectoderm would result in enhancement of SAC efficiency. In *C. intestinalis* the transcription factor FoxA-a is necessary for the acquisition of anterior ectodermal identity and its overexpression in animal blastomeres from the 16-cell stage results in posterior cells acquiring anterior identity (Lamy et al., 2006). To induce anterior fate in the posterior part of the ventral ectoderm we injected *P. mammillata* eggs with a plasmid encoding FoxA-a under the control of the Fog promoter (pFog>FoxA-a) to drive FoxA-a expression in all animal cells (Lamy et al., 2006). At the 9^th^ cell cycle, in pFog>FoxA-a injected embryos expression of the anterior marker *sFRP1/5*, normally restricted to the anterior pole (Fig. S3C), was extended towards the posterior pole in 34% of embryos and was expressed in all cells in 66% of embryos. The latter did not elongate at the neurula stage but developed into round embryos, indicating complete loss of anterior-posterior patterning. At the 9^th^ cell cycle, mitotic duration was unaffected in pFog>FoxA-a-injected embryos in the presence of DMSO (Fig. S3D). In nocodazole-treated round pFog>FoxA-a-injected embryos (Fig. 3G) mitotic duration was extended by 4.4-fold (11.1±2.6 to 48±21.7 minutes), an increase similar to that observed in anterior wild-type cells (9.9±2.4 to 49.3±11.9 minutes). To confirm that anteriorization of the posterior ectoderm results in an increase in SAC strength, we also removed the vegetal pole of fertilized eggs, which contains the maternal factors required for cell differentiation and for the invariant cleavage pattern typical of ascidian development (Fig. 4A). Such manipulation of the zygote (pole-ablated) gives rise to embryos in which all cells acquire the anterior ectodermal fate (Dumollard et al., 2017; Nishida, 1996), as shown by ectopic expression of the anterior marker *sFRP1/5* throughout the embryo (Fig. S3C). Pole-ablated embryos acquire SAC activity at the 8^th^ cell cycle and in the presence of nocodazole delay mitotic duration by 2.4-fold (DMSO: 10.7±2.9 to nocodazole: 24.3±7.5 minutes), like ventral ectodermal cells in un-manipulated embryos (Fig. S3E). At the 9^th^ cell cycle, we observed greater variability in mitotic duration in pole-ablated embryos than in anterior ectodermal cells of wild-type embryos, with several cells exhibiting only a short mitotic delay (Fig. 4B). However, we noticed that differently from unmanipulated control embryos where the cell volume of anterior ectodermal cells ranges between 2×10^3^ µm^3^ and 4×10^3^ *µ*m^3^, in pole-ablated embryos cell volume varied between 1.8×10^3^ *µ*m^3^ and 8×10^3^ *µ*m^3^ (Fig. 4C). The mitotic delay induced by nocodazole was comparable in control (48±11.9 minutes) and pole ablated (43.5±15.6 minutes) embryos when considering only cells whose volume was comparable (volume <4×10^3^ *µ*m^3^), while in pole ablated cells larger than 4×10^3^ *µ*m^3^ the mitotic delay was significantly shorter (25.3±6.9 minutes, 2.2-fold increase; Fig. 4C, D). Hence, the difference in SAC efficiency between anterior and posterior ectodermal cells could be attributed to a difference in cell identity. Our data also suggest that within same fated cells, the SAC response is modulated by cell size and is stronger in smaller cells.

**Figure 4:**
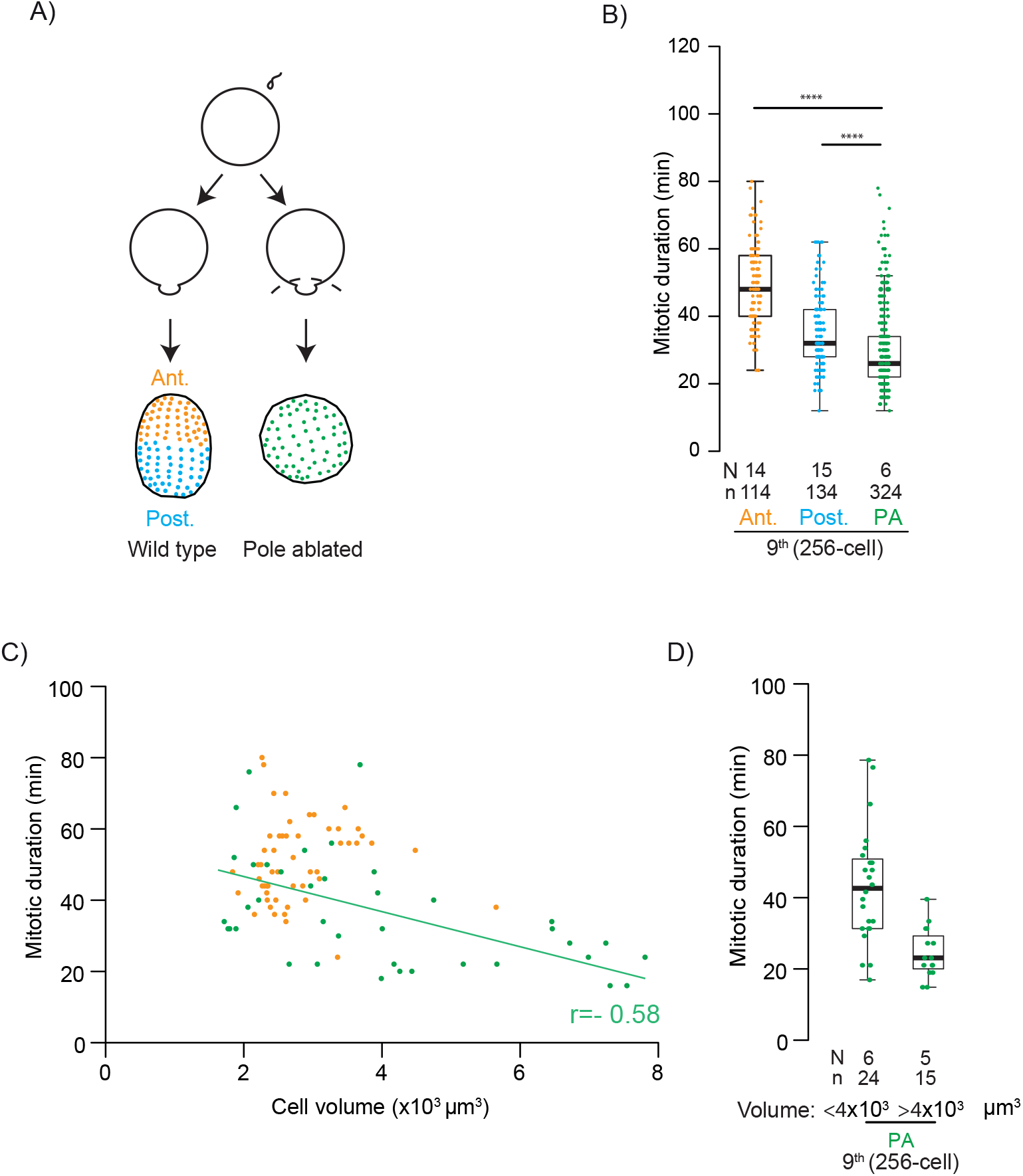
SAC strength depends on cell size. **A)** Schematic representation of non-manipulated (left) and pole-ablated (right) fertilized eggs and resultant embryos. **B)** Mitotic duration in the presence of nocodazole for wild type ventral anterior (orange) and posterior (blue) ectodermal cells and cells from pole-ablated embryos (green) at the 9^th^ cell cycle. **C)** Mitotic duration in pole-ablated embryos and anterior ectodermal cells treated with nocodazole as a function of cell volume at the 9^th^ cell cycle. The line indicates the regression fit for pole ablated cells and r the coefficient of correlation for the two variables in the same cells. **D)** Mitotic duration in small (< 4×10^3^ *µ*m^3^) and large (> 4×10^3^ *µ*m^3^) cells from nocodazole treated pole-ablated (green) embryos at the 9^th^ cell cycle. For B), C) and D) boxes are as in Figure 1B.

Thus, following SAC acquisition, SAC strength is specific to different cell populations and is modulated by cell size. This is similar to what was reported for early *C. elegans* embryos, where SAC strength increases at each cleavage division (Galli and Morgan, 2016), in a lineage specific manner (Gerhold et al., 2018). Modulation of SAC strength in SAC proficient cells, therefore, appears to follow similar rules in both chordate and non-chordate embryos. Our results also suggest that GSK3 is involved in enhancing SAC efficiency in anterior ectodermal cells. GSK3 activity as a SAC enhancing factor could be direct through phosphorylation of the SAC machinery and/or the kinetochores to favor their interaction, or indirect through ß-catenin mediated transcriptional control of SAC regulators. Although further work will be required to distinguish between these possibilities, a study carried out in mammalian cells showed that GSK3 promotes the recruitment of the SAC components Mad2, Bub1 and BubR1 to unattached kinetochores and favors the formation of the inhibitory complex (Rashid et al., 2018), supporting a direct role of GSK3 in increasing SAC strength.

## Material and Methods

### Gamete preparation

All experiments were performed using the ascidian *Phallusia mammillata*, a marine chordate. Animals were collected at Sète (France) and maintained in aquaria at the Laboratoire de Biologie du Developpment de Villefranche sur mer (LBDV).

Dissection of adult animals allowed collection of both sperm and eggs. Dry sperm was kept at 4°C for a maximum of 2 weeks. Eggs were collected in microfiltered sea water (MFSW) containing 5 mM TAPS ([tris(hydroxymethyl)methylamino]propanesulfonic acid), pH 8.2 (McDougall et al., 2015). The chorion was removed by treatment with 0.1% trypsin in 10 ml MFSW-TAPS for 1.5-2 hours at room temperature (Sardet et al., 1989). Dechorionated eggs were washed in MFSW and kept at 18°C in MFSW with TAPS. Eggs were used within 36 hours from collection. All glass- and plastic-ware for embryo handling was coated with 0.1% gelatin and 0.1% formaldehyde (GF) (McDougall et al., 2015).

Prior to fertilization sperm was activated by incubating 9 *µ*l of sperm (from 3 different animals) in 1 ml of MFSW pH 9.2 for 30 minutes. Activated sperm was added to the eggs at a ratio of approximately 10 *µ*l of sperm to 5-20 eggs for small cultures or injected eggs, and 100 *µ*l for 10 ml of MFSW for large cultures. Following fertilization eggs were transferred to a fresh dish to remove excess sperm.

### Plasmids and RNAs

Plasmids encoding NLS-3Venus, NLS-tomato, PH-tomato and PH-GFP were previously described (McDougall et al., 2015). Following plasmid linearization with Sfi1, RNAs were synthetized using T3 RNA polymerase (mMessage-mMachine kit, Invitrogen). NLS-3Venus and NLS-tomato RNAs were injected in unfertilized eggs at the concentration of 4 *µ*g/*µ*l. PH-tomato and PH-GFP were injected in unfertilized eggs at a concentration of 1 *µ*g/*µ*l. At these concentrations, these RNAs did not delay mitotic progression and had no effect on embryonic development.

The cDNA encoding Mad2-DN, a phosphomimic form of the human Mad2 that acts as a dominant negative, was provided by Dr K. Wassmann (Wassmann et al., 2003a) and was cloned into the plasmid pCS2 to allow RNA synthesis using SP6 RNA polymerase (mMessage-mMachine kit, Invitrogen, Carlsbad, USA). Following in vitro transcription, a poly(A) tail was added using a polyA-tailing kit (Invitrogen). The Mad2-DN RNA was injected into unfertilized eggs at a concentration of 8 *µ*g/*µ*l.

FoxA-a identified as Phmamm.g00001891 (Brozovic et al., 2018) was amplified from a cDNA template using the primers: forward cgcTGTACAAGATGATGTTGTCGTCTCCC and reverse cgcTGTACATTAGTTGGCCGGTACGCA. The FoxA-a coding sequence was used to replace the RFA cassette in pSP72BSSPE-pFog::Venus-RFA provided by Patrick Lemaire (Roure et al., 2007) by enzymatic digestion using BsrGI. The pFog>FoxA-a plasmid was injected at a concentration of 35 ng/*µ*l.

### Microinjection

Microinjection of unfertilized eggs was carried out as previously described (Yasuo and McDougall, 2018). Briefly glass capillaries without filament (GC100 T10, Harvard Apparatus) were used to store the RNA to be injected. Prior to loading, the RNA solution was centrifuged at 13000 rpm at 10°C for 10 min to pellet impurities that could block the needle. Capillaries (GC100T10 Harvard Apparatus) were cut to 1cm length and filled sequentially with 2 *µ*l of mineral oil, 0.5-1.5 *µ*l RNA or DNA solution and finally 1 *µ*l of mineral oil and kept at 4°C for several weeks.

Injections were performed using a Leica DM IL LED inverted microscope (10x objective) in transmitted light. Needles were made by pulling capillaries with a Narishige horizontal puller (PN-30). The needle was attached to a NarishigeIM300 injection box and manipulated using a three-axis hydraulic micromanipulator (Narishige MMO-203 associated with NarishigeNO-SIX-2 mounting adaptor and Narishige HI-7 pipette holder). Eggs were immobilized in horizontal injection chambers (Yasuo and McDougall, 2018) filled with MFSW and injected to 5% of their volume.

### Pole ablation

Following fertilization, the cytoplasm of ascidian embryos undergoes significant rearrangements that lead to the localization of many maternal factors into a transient surface protrusion known as the contraction pole (Nishida, 1994; Nishida, 1996). Ablation of this pole was performed by aspiration with a micropipette as previously described (Nishida, 1996). An aspiration micropipette, formed by breaking the tip of a microinjection needle to obtain a blunt opening, was connected to a mouth pipette. Eggs were fertilized in a petri dish and immediately transferred into an injection chamber mounted on an inverted microscope as for microinjection. The contraction pole was aspirated into the pipette by suction and a quick movement of the needle perpendicularly to the egg surface sealed the membrane of the zygote. Eggs were then transferred to a new dish to allow development.

### Drug treatments

All drugs were stored in small aliquots at -20°C.

Nocodazole (Sigma-Aldrich, St. Louis, MO, USA) was resuspended in DMSO to obtain a 33 mM stock solution and was used at a final concentration of 10 *µ*M in MFSW. Reversine (Axon Medchem, Groningen, The Netherlands) was resuspended in DMSO at 5 mM and used at a final concentration of 0.5 *µ*M in MFSW. Both drugs were added to MFSW when embryos reached the analyzed stage and embryos were then maintained in the presence of the drug for the duration of the experiment.

BIO ((2’Z,3’E)-6-Bromoindirubin-3’oxime, Millipore) was resuspended in DMSO to obtain a 10 mM stock solution and was then used at a final concentration of 2.5 *µ*M in MFSW. Embryos were soaked in 2.5 *µ*M BIO, in the dark, from the onset of gastrulation (stage 10, about 5hpf). Two hours later (about 7 hpf) embryos were treated with 10*µ*M nocodazole and maintained in the presence of both drugs for the remaining of the experiment.

### Immunofluorescence

For immunofluorescence, embryos were collected at the required stage and fixed overnight in 100% methanol at -20°C. For Mad1 staining, embryos were treated with 10*µ*M nocodazole before fixation. After fixation, embryos were washed 3 times in PBS containing 0.1% triton-X-100 (PBST), then incubated in PBST containing 3% BSA (PBST-BSA) for 1 hour at room temperature (RT) and finally incubated overnight at 4°C in PBST-BSA with the anti-tubulin antibody, DM1A (mouse, Sigma-Aldrich) at 1:500 dilution or a specific antibody raised against *P. mammillata* Mad1 at 1:100 dilution (Chenevert et al., 2020). After overnight incubation, excess antibody was removed by 3 consecutive washes in PBST. Embryos were then incubated in PBST-BSA with either Alexa fluor 546 or Alexa fluor 488 anti-mouse secondary antibody (Jackson ImmunoResearch, Ely, UK; 1:1000) at RT for 1-2 hours. Following 2 further washes in PBST, embryos were incubated for 10 minutes in PBST containing Hoechst 33342 (5 *µ*g/ml), washed twice and then mounted in citifluor AF1 (Science Services, München, Germany), for imaging with a Leica SP8 inverted confocal microscope using a 40x water objective.

### Time-Lapse Microscopy

For live imaging, embryos were mounted between GF coated slide and coverslip, using Dow Corning vacuum grease as spacer. In early stages (2- to 16-cell embryos) nuclei were followed by brightfield microscopy while at later stages (32-512 cell) we relied on the localization of a fluorescent protein, Venus or Tomato, fused to a nuclear localization signal (NLS-3Venus or NLS-Tomato). The NLS protein accumulates in the nucleus during interphase, while from NEB to NER (mitosis) the signal disperses in the cytoplasm. To distinguish individual cells within the embryo, eggs were co-injected with an mRNA encoding the plasma-membrane marker PH domain-GFP. Following fertilization embryos were maintained in MFSW until they reached the stage of interest and then treated with the different drugs (nocodazole, reversine, BIO) or DMSO and transferred to the appropriate microscope for recording. Live imaging was always performed in a temperature-controlled room set at 19°C. When possible, control and treated embryos were imaged in parallel on the same microscope.

Image acquisition was performed on the Plateforme d’Imagerie Microscopique de Villefranche (PIM). For brightfield imaging, a z-stack covering the entire embryo with a z step of 2 *µ*m was acquired using a Zeiss (Oberkochen, Germany) Axiovert 100 or Axiovert 200 inverted microscope with bright field optics fitted with a 40x/1.1NA objective and equipped with a CoolSnap Kino camera (Photometrics) and Metamorph acquisition software. Images were acquired every 1-2 minutes. For NLS expressing embryos, a z-stack covering the entire embryo with a z step of 2 *µ*m was acquired using a Leica (Wetzlar, Germany) SP8 inverted confocal microscope fitted with a 40x water immersion objective and 408, 488, and 552 nm lasers and the Leica application suite acquisition software (LASX). Images were acquired every 2 minutes.

### Image analysis

Images were analyzed using different softwares: Metamorph (Molecular Devices, San Jose, USA), imageJ-Fiji (Schindelin et al., 2012), Imaris (Imaris, BitPlane, Zurich, Switzerland) and Icy (de Chaumont et al., 2012) with the plugin Z-explorer (https://icy.bioimageanalysis.org/plugin/z-explorer/). Presence or absence of nuclei was determined manually by following each cell over time through the entire Z stack. At the 512-cell stage, if a cell did not exit mitosis for more than 1 hour before the end of acquisition, we assigned a mitotic duration of 60 minutes. Approximate cell number was assessed by counting number of nuclei using the Spot tool in Imaris. Cell volume was determined using the Surface tool in Imaris: the cell contour was drawn in each frame based on the signal from the plasma-membrane associated PH-GFP or PH-Tomato, the reconstructed cell shape was confirmed using 3D visualization of the embryo and then the cell volume was retrieved.

### Western blot

To assess Mad1 levels, 30 eggs or embryos were collected for each stage from egg to 512-cell stage. Embryos were collected in MFSW and mixed with the same volume of 2x Laemmli (100 mM Tris-HCl at pH 6.8, 4% SDS, 0.2 % bromophenol blue, 20 % glycerol, 200 mM dithiothreitol). Samples were heated for 5 minutes at 95°C and then kept at -20°C. Samples were separated on a 10% SDS-polyacrylamide gel and then transferred to a nitrocellulose membrane. Membranes were blocked for 1h in Tris Buffer-Saline (TBS: 20mM Tris Base, 150 mM NaCl) containing 2% milk, 0.1% tween-20 (TBS-Tween-milk) and then incubated in blocking solution containing the appropriate antibody overnight at 4°C. For Mad1 detection the Mad1 antibody (Chenevert et al., 2020) was diluted 1:2000, whereas for tubulin detection, the DM1a antibody (Sigma-Aldrich, mouse) was diluted 1:3000. Following antibody incubation, membranes were washed 3 times with TBS-Tween (TBS with 0.1% tween-20) and incubated with an appropriate horseradish peroxidase-conjugated secondary antibody (Jackson ImmunoResearch) at 1:10000 dilution in TBS-Tween-milk for 1 hour at RT. After 3 washes with TBS-Tween, signal detection was carried out using the SuperSignal West Pico chemio-luminescent substrate (Thermo Scientific, Waltham, MA, USA) as described by the manufacturer and imaged in the Vilmen-Loubet Fusion machine.

### *In situ* hybridization

The *sFRP1/5* sequence identified as Phmamm.g0000599 (Brozovic et al., 2018) and the *FoxF* sequence identified as Phmamm.g07919 were amplified from cDNA templates. After linearization by enzymatic digestion with SalI, an *in situ* hybridization probe covering the coding sequence, was generated by in vitro transcription using T7 RNA polymerase (Promega, Charbonnières-les-Bains, France) in presence of Digoxygenin-labeled nucleotides (DIG RNA Labeling Mix, Roche, Bâle, Switzerland).

*In situ* hybridization experiments were performed as previously described (Paix et al., 2009). Eggs and embryos were fixed overnight at 4°C in 4% formaldehyde, 100 mM MOPS, 0.5 M NaCl, pH 7.6, washed 3 times in PBS, progressively dehydrated in ethanol (25%, 50%, 75%, 100%) and stored at −20°C. Fixed eggs and embryos were then re-hydrated by addition of an equal volume of PBS/0.1% Tween 20 (PBSTw), washed once in PBSTw and then incubated with 2 μg/ml proteinase K (Sigma) in PBSTw for 25 minutes at room temperature (RT). After 3 washes in PBSTw, embryos were fixed again in PBS containing 4% formaldehyde (1 h, RT) and washed again to remove formaldehyde. Embryos were then incubated in hybridization buffer (50% formamide, 6× saline sodium citrate (SSC), 5× Denhardt’s solution, 1 mg/ml yeast RNA, 0.1% Tween, pH = 7.5) for 1 h at 65 °C. Probes were added to fresh hybridization solution at a final concentration of 0.5 ng/μl and hybridization was performed overnight at 65°C. After hybridization, embryos were washed at 65°C twice in 50% formamide, 5× SSC, 1% SDS (30 min), twice in 50% formamide, 2× SSC, 1% SDS (15 min), once in 2× SSC, 0.1% Tween (15 min) and twice in 0.2× SSC, 0.1% Tween (15 min). All solutions were at pH 7.5. Embryos were then incubated in blocking buffer (0.15 M NaCl, 0.5% BBR, 0.1M Tris pH8) for 1h at RT. Anti-DIG AP antibody (Roche, Bâle, Switzerland, 1:4000) was added in fresh blocking solution and incubated at 4 °C overnight. After incubation, embryos were washed 5 times in PBSTw (3 times for 10 min, 30 min, 1 h respectively) and then in TBSTw at 4°C overnight. Embryos were washed once with reaction buffer (0.1 M Tris pH 9.5, 50 mM MgCl_2_, 0.1M NaCl, 0.1% Tween) and then the detection reaction was carried out in a buffer containing 3.5 *µ*l/ml of of nitro blue tetrazolium (NBT) and 5-bromo-4-chloro-3-indolyl-phosphate (BCIP) (Roche, Bâle, Switzerland). The reaction was stopped with 50 mM EDTA in PBS and embryos were washed in PBSTw. Embryos were fixed in 4% formaldehyde in PBS, washed in PBS and mounted in 80% Glycerol. Stained embryos were imaged using an optical microscope (Zeiss Imager A2) equipped with a xxx camera and Zeiss Zen blue software, using a 20x objective.

### Quantification and statistical analysis

Number of analyzed embryos (N) and cells (n) are reported under each graph. For each experiment at least 3 independent biological repeats were included. For statistical analysis the non-parametric test of Wilcoxon-Mann-Whitney was applied. For all comparisons, we used population medians as the mean can be affected by extreme values and is highly impacted by outliers (Gaddis and Gaddis, 1990). However, means and standard deviations are also provided in the text. Statistical tests and graphics were performed using the R software (R Core Team, 2016) and the library “pretty R” (Grosjean and Lemon, 2015) was used to retrieve median, mean and standard deviation.

Box plot represents the distribution of the data using quartiles (Q). Q1, Q2 and Q3 being respectively the values below which lie 25%, 50% and 75% of the data points. Q1 and Q3 form the limits of the box and the median Q2 is indicated by a bolder line. The whiskers represent the range in which most values are found, whereas the values outside represent the outliers.

## Supporting information

Movie 1: P. mammillata 8-cell stage embryos undergoing mitosis with and without spindle microtubules.

Movie 2: P. mammillata 256-cell stage embryos undergoing mitosis with and without spindle microtubules.

Movie 3: SAC-impaired P. mammillata 2-cell stage embryos undergoing mitosis with and without spindle microtubules.

Movie 4: SAC-impaired P. mammillata 256-cell stage embryos undergoing mitosis with and without spindle microtubules.

## Acknowledgments

We are grateful to E. Houliston, H. Yasuo, C. Hudson, S. Darras, R. Dumollard, P. Lemaire and K. Wassmann for discussion, sharing of reagents or critical reading of the manuscript. We would also like to thank C. Hebras, F. Bekkouche, L. Gilletta, M. Temagoult, S. Benaicha, and S. Schaub for technical assistance. We thank the Service Aquarologie of CRB (IMEV-FR3761) for access to marine facilities and animals and the microscopy platform PIM (member of the MICA microscopy platform) where image acquisition was conducted. CRB and PIM are supported by EMBRC-France, whose French state funds are managed by the ANR within the investments for the future program under reference ANR-10-INSB-02.

This research was funded by the CNRS and from PACA region (project Mepanep, 2014_03738).

