## Supplemental Figure 1 for "Acquisition of the Spindle Assembly Checkpoint and its modulation by cell fate and cell size in a chordate embryo"

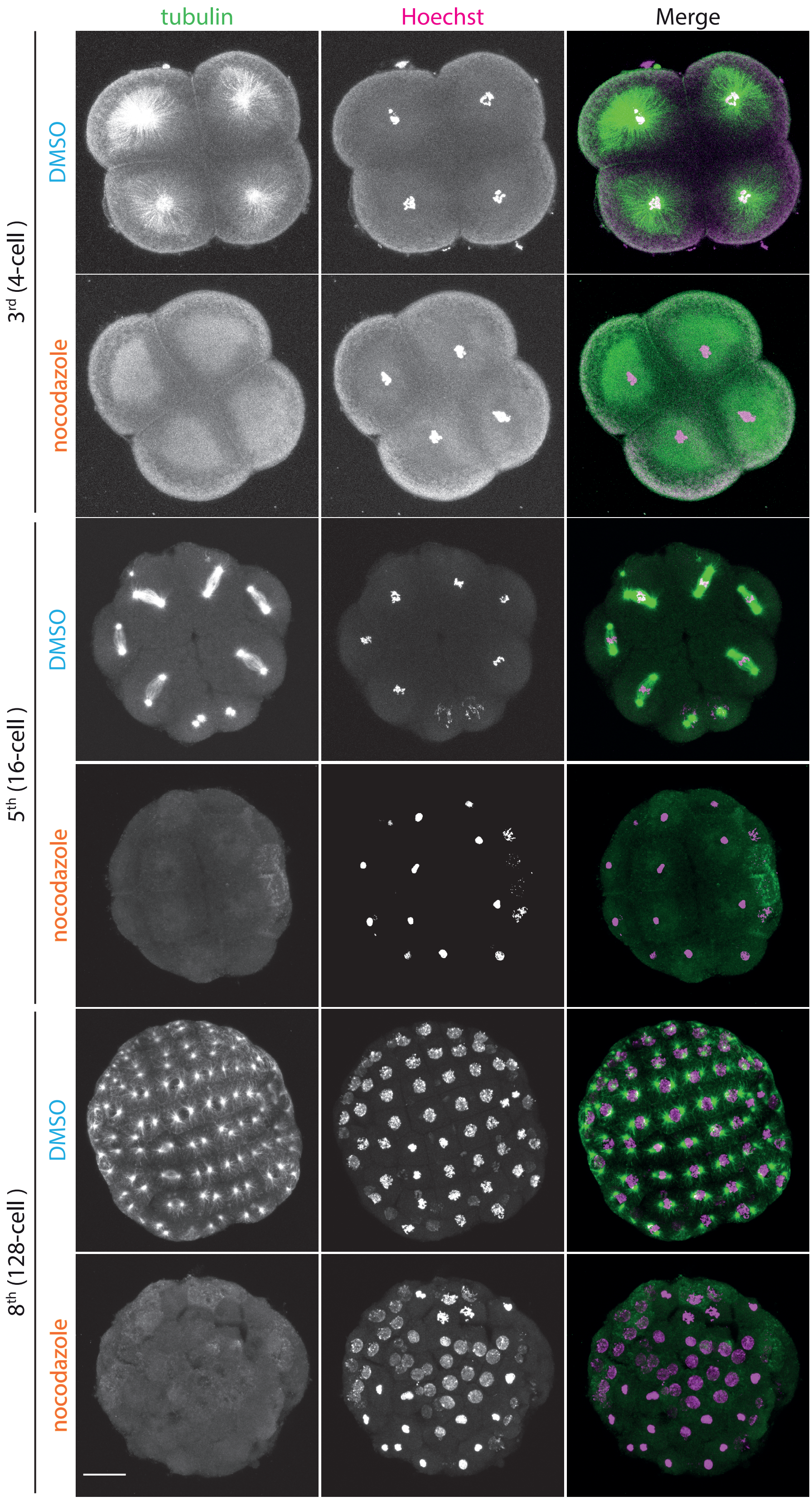


**Figure S1: Nocodazole induces microtubule depolymerization in *P. mammillata* embryos.**

Representative images of 4-cell, 16-cell and 128-cell embryos stained for microtubules (anti-tubulin, green) and for DNA (Hoechst, magenta) following treatment with either DMSO or 10 $\mu$M nocodazole. Images are Z-projections of confocal stacks spanning the whole embryo. Scale bar is 30 $\mu$m.
