## Supplemental Figure 2 for "Acquisition of the Spindle Assembly Checkpoint and its modulation by cell fate and cell size in a chordate embryo"

A)

```
Hs_Mad2    MALQLSREQGITLRGSAEIVAEFFSFGINS 30
Pm_Mad2    -MAAAKQLNKITLKGSAAIVSEFFFYGMNN 29
                  .: :  ***:***  **:***  :*:*

Hs_Mad2    ILYQRGIYPSETFTRVQKYGLTLLVTTDLE 60
Pm_Mad2    ILYQRGVYPPEMFKQEKKYGLTILVSTDET 59
           .******:** * *.:  :*****:**:**

Hs_Mad2    LIKYLN-NVVEQLKDWLYKCSVQKLVVVIS 89
Pm_Mad2    LLTYLNDNVLPQLQEWLEKGAVKGLVVVMR 89
           *:.*** **:  **::** *  :*:  ****:

Hs_Mad2    NIESGEVLERWQFDIECDKTAKDDSAPREK 119
Pm_Mad2    ESETNETLEGWQFEIQCESDKSENGKPATK 119
           : *:.*.** ***:*:*:.   .::. *  *

Hs_Mad2    SQKAIQDEIRSVIRQITATVTFLPLLEVSC 149
Pm_Mad2    SKETINSEIRAVIRQVTATVTFLPLLECPC 149
           *:::*:.***:****:***********  *

Hs_Mad2    SFDLLIYTDKDLVVPEKWEESGPQFITNSE 179
Pm_Mad2    SFDLLFYTDKNLEVSEKWEDSSPCLMPNSE 179
           *****:****:* * ****:*.*  :: ***

Hs_Mad2    EVRLRSFTTTIHKVNSMVAYKIPVND 205
Pm_Mad2    EVKLGSFSTSVHKVDGAVSYKFSM-- 203
           **:* **:*::***:. *:**: :
```

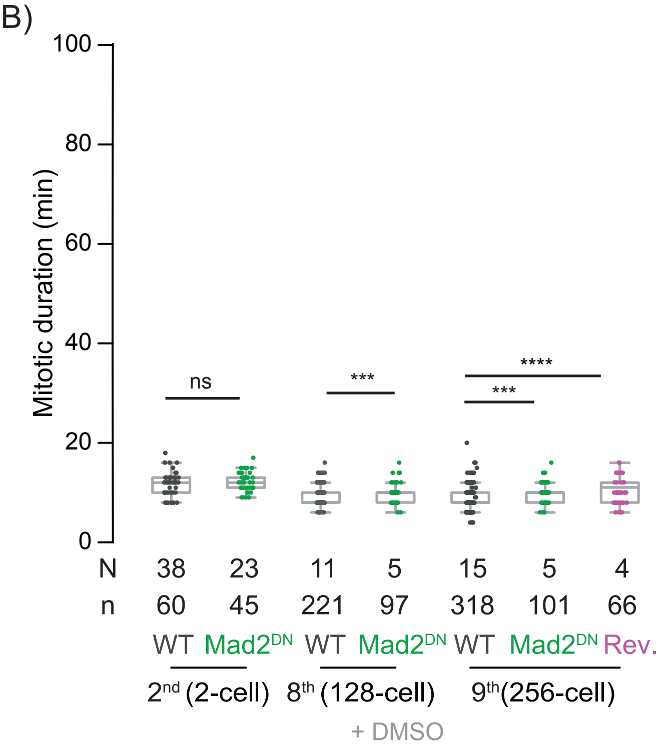


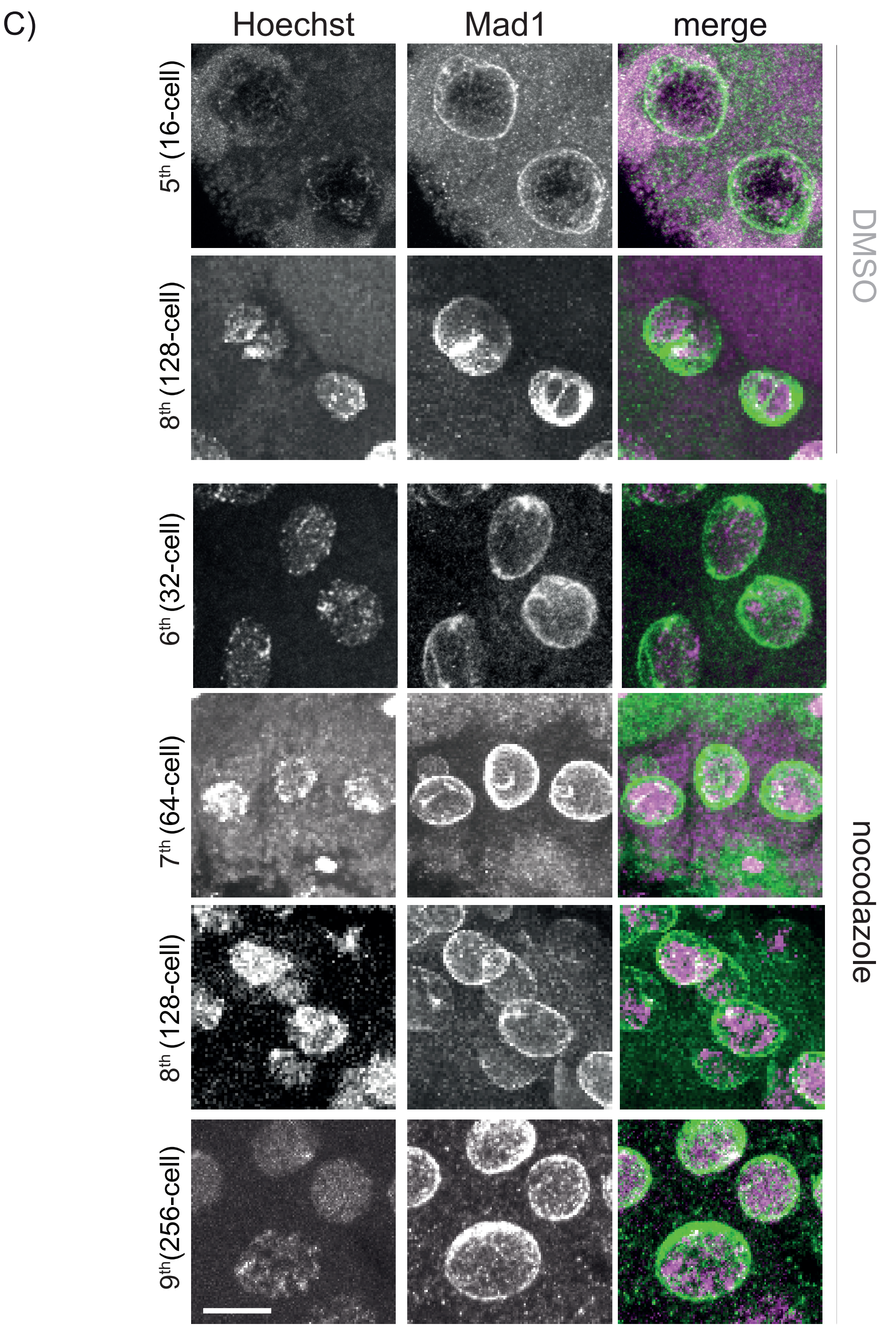


Figure S2

**Figure S2: Mad2 sequence alignment and Mad1 protein localization during interphase.**

**A)** Alignment of Mad2 sequences from human and from *P. mammillata*. Phosphorylation sites mutated in the Mad2-DN construct are marked in red. **B)** Quantification of mitotic duration in DMSO- treated embryos at 2-, 128- and 256-cell stage in wild-type embryos (black) or following SAC impairment by either overexpression of Mad2-DN (green) or treatment with reversine (magenta). Drugs (nocodazole and reversine) were added when embryos reached the analyzed stage. Boxes are as in Figure 1B. **C)** Representative images of interphase cells from DMSO treated 16- and 128-cell stage embryos and nocodazole treated embryos at 32-, 64-, 128- and 256-cell stage, fixed and stained for Mad1 (green) and Hoechst (DNA, magenta). Scale bar is 10 $\mu$m.
