## Supplemental Figure 3 for "Acquisition of the Spindle Assembly Checkpoint and its modulation by cell fate and cell size in a chordate embryo"

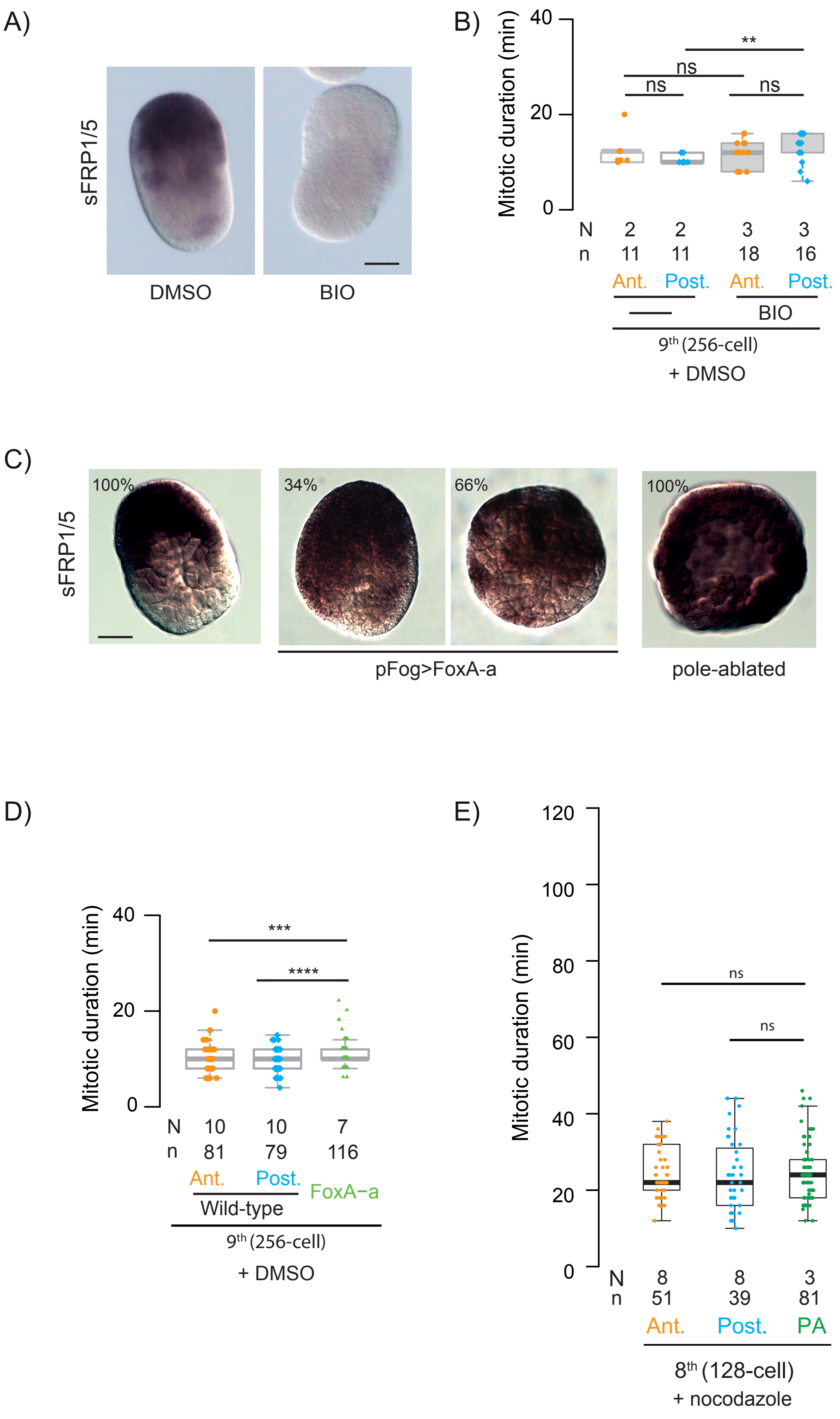
A)
sFRP1/5
DMSO
BIO
B)
Mitotic duration (min)
40
20
0
**
ns
ns
ns
N 2 2 3 3
n 11 11 18 16
Ant. Post. Ant. Post.
BIO
9th (256-cell)
+ DMSO
C)
sFRP1/5
100%
34%
66%
100%
pFog>FoxA-a
pole-ablated
D)
Mitotic duration (min)
40
20
0
***
****
N 10 10 7
n 81 79 116
Ant. Post.
Wild-type
FoxA−a
9th (256-cell)
+ DMSO
E)
Mitotic duration (min)
120
100
80
60
40
20
0
ns
ns
N 8 8 3
n 51 39 81
Ant. Post. PA
8th (128-cell)
+ nocodazole

Figure S3

**Figure S3: Anterior-posterior polarity in BIO-treated embryos and pole-ablated embryos**

**A)** *In situ* hybridization showing the expression pattern of the anterior marker *sFRP1/5* in un-manipulated (left) and Bio-treated (right) embryos. **B)** Mitotic duration in anterior (orange) and posterior (blue) ventral ectodermal cells of control (left) and BIO-treated (right) embryos at the 9$^{th}$ cell cycle in DMSO. **C)** *In situ* hybridization showing the expression pattern of the anterior marker *sFRP1/5* in un-manipulated (left), pFog>FoxA-a expressing (middle; n=64, 22 neurula-like and 42 round embryos) and pole-ablated (right) embryos. **D**) Mitotic duration in wild type anterior (orange) and posterior (blue) ventral ectodermal cells and pFog>FoxA-a over-expressing (green) embryos at the 9$^{th}$ cell cycle in DMSO. **E)** Mitotic duration in nocodazole treated anterior (orange) and posterior (blue) cells in wild type and in pole-ablated (green) embryos at the 8$^{th}$ cell cycle. Boxes are as in Figure 1B. Scale bars are 30 $\mu$m.
